## Supporting Information for "Tackling Recalcitrant *Pseudomonas aeruginosa* Infections In Critical Illness via Anti-virulence Monotherapy"

T. Kitao, Department of Microbiology, Graduate School of Medicine, Gifu University, Gifu 501-1194, Japan

A Felici, Academic Partnership, Evotec SE, 37135 Via A. Fleming 4, Verona, Italy

**Supplementary Table 1A-G. Structure activity relationship (SAR) studies.**
The assessment of inhibitors potency was based on pyocyanin production, *pqs* operon gene expression using the *pqsA*-GFP reporter, and the production of the MvfR-regulated low molecular weight molecules HHQ, PQS, HQNO, DHQ, 2-AA in the presence of 50µM of each inhibitor. Assessment of the anthranilic acid (AA) production was used as additional control of the inhibition of the low molecular weight molecules since their synthesis depends on AA, their primary precursor.

**A): Optimization starting with M17**

SAR studies of N 2-[(4-fluoro-2-methylphenyl)amino]-N-phenylacetamide as anti-MvfR agents, via exploration of the aryl portion of the N-phenylacetamide side.

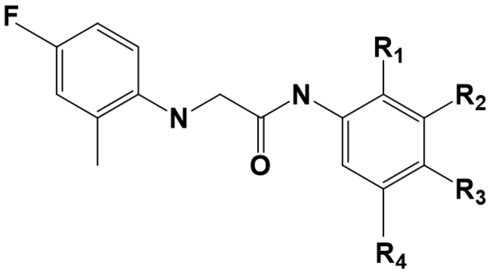


| Compound | | | | | % Production | | | | | | | |
|---|---|---|---|---|---|---|---|---|---|---|---|---|
| | R1 | R2 | R3 | R4 | Pyocyanin | *pqsA*-GFP | HHQ | PQS | HQNO | 2-AA | DHQ | AA |
| **M17** | | | **CN** | | **67** | **32** | **3** | **11** | **17** | **101** | **135** | **100** |
| G1 | | CN | | | 103 | 84 | 100 | 109 | 101 | 122 | - | - |
| G2 | | | $CF_3$ | | 61 | 46 | 110 | 93 | 86 | 68 | 81 | - |
| G3 | CN | | | | 113 | 96 | 99 | 122 | 111 | 123 | - | - |
| G4 | | | F | | 84 | 92 | 95 | 88 | 92 | 80 | 85 | - |
| D2 | Br | | | | 101 | 96 | - | - | - | - | - | - |
| D5 | | | 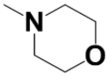 | | 102 | 100 | - | - | - | - | - | - |
| D7 | Me | | Br | | 89 | 70 | - | - | - | - | - | - |
| D9 | | | 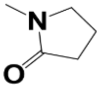 | | 107 | 105 | - | - | - | - | - | - |
| D10 | F | | | F | 90 | 89 | - | - | - | - | - | - |
| D12 | Br | | $CH_3$ | | 108 | 92 | - | - | - | - | - | - |
| D13 | | CN | | | 100 | 90 | - | - | - | - | - | - |
| D14 | | $NO_2$ | | | 85 | 77 | - | - | - | - | - | - |
| D15 | | | OMe | | 102 | 87 | - | - | - | - | - | - |
| D19 | | | 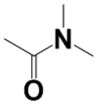 | | 109 | 100 | - | - | - | - | - | - |
| D21 | Me | | Me | | 113 | 87 | - | - | - | - | - | - |
| D22 | | Cl | | | 96 | 71 | - | - | - | - | - | - |
| D30 | 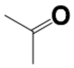 | | | | 91 | 94 | - | - | - | - | - | - |
| D31 | | $CH_3$ | Br | | 69 | 47 | - | - | - | - | - | - |
| D32 | | OMe | | | 92 | 94 | - | - | - | - | - | - |

**B.** SAR studies of N-(4-cyanophenyl)-2-(phenylamino) acetamide as anti-MvfR agents via exploration of the aryl portion of the 2-phenylamino portion of the molecule.

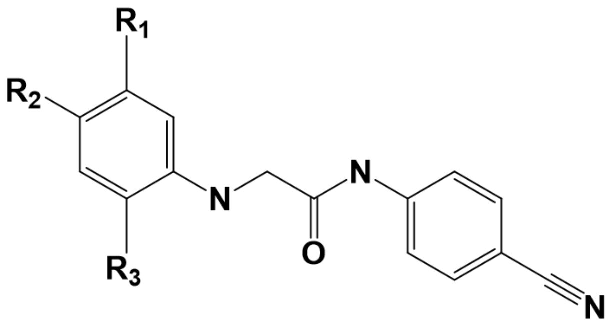

| Compound | | | | % Production | | | | | | | |
|---|---|---|---|---|---|---|---|---|---|---|---|
| | **$R_1$** | **$R_2$** | **$R_3$** | **Pyocyanin** | **pqsA -GFP** | **HHQ** | **PQS** | **HQNO** | **2-AA** | **DHQ** | **AA** |
| G8 | | F | | 86 | 63 | 87 | 124 | 108 | 112 | - | - |
| G9 | | | Me | 94 | 93 | 76 | 124 | 92 | 64 | 59 | - |
| D3 | | | Et | 97 | 92 | - | - | - | - | - | - |
| D6 | F | | | 91 | 68 | - | - | - | - | - | - |
| D16 | | Cl | | 43 | 18 | - | - | - | - | - | - |
| D18 | F | | Me | 69 | 28 | - | - | - | - | - | - |
| D25 | | | F | 97 | 84 | - | - | - | - | - | - |
| D28 | F | $CH_3$ | | 60 | 42 | - | - | - | - | - | - |
| D33 | | $CH(CH_3)_2$ | | 59 | 53 | - | - | - | - | - | - |

**C**. SAR studies retaining the 4-cyanophenyl and exploring the central linker and the effect of the substituent on the second Aryl group.

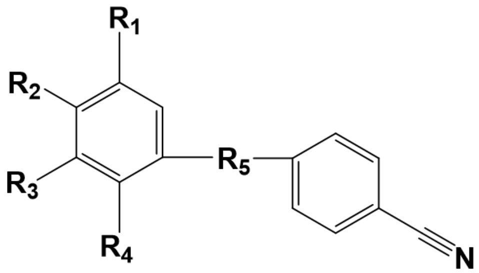


| Compound | | | | % Production | | | | | | | |
|---|---|---|---|---|---|---|---|---|---|---|---|
| | $R_2$ | $R_4$ | $R_5$ | Pyocyanin | *pqsA* -GFP | HHQ | PQS | HQNO | 2-AA | DHQ | AA |
| G5 | F | Me | 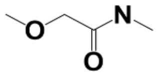 | 98 | 98 | 114 | 114 | 111 | 63 | 83 | 54 |
| G6 | F | Me | 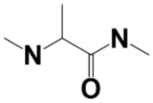 | 110 | 87 | 106 | 115 | 131 | 152 | - | - |
| D38 | F | Me | 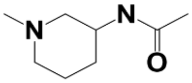 | 108 | 90 | 98 | 96 | 100 | 101 | 84 | 54 |
| D39 | F | Me | 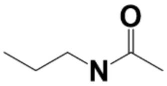 | 110 | 91 | 96 | 93 | 102 | 106 | 96 | 68 |
| D40 | F | Me | 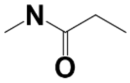 | 115 | 96 | 102 | 100 | 109 | 113 | 85 | 81 |
| D24 | F | | 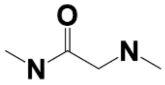 | 59 | 26 | - | - | - | - | - | - |
| D34 | F | | 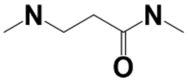 | 91 | 80 | 97 | 97 | 96 | 113 | 100 | 101 |
| D37 | F | | 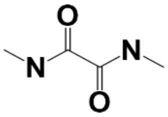 | 106 | 97 | 98 | 110 | 99 | 103 | 105 | 90 |
| D36 | F | | 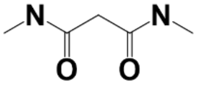 | 16 | 10 | 57 | 47 | 59 | 37 | 68 | 187 |
| D44 | F | | 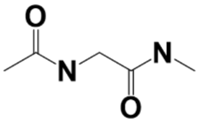 | 94 | 71 | 94 | 92 | 94 | 92 | 97 | 32 |
| D47 | F | | 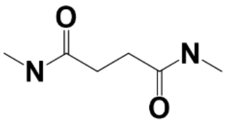 | 99 | 78 | 95 | 94 | 93 | 100 | 100 | 42 |
| D83 | CN | | 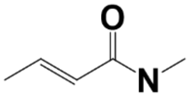 | 114 | - | - | - | - | - | - | - |

| | | | | | | | | | | |
|---|---|---|---|---|---|---|---|---|---|---|
| D76 | F | 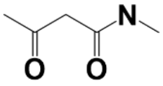 | 77 | - | - | - | - | - | - | - |
| D84 | F | 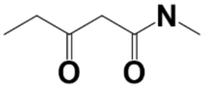 | 102 | - | - | - | - | - | - | - |
| D86 | F | 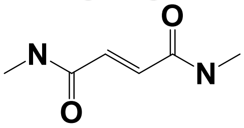 | 88 | - | - | - | - | - | - | - |
| D89 | F | 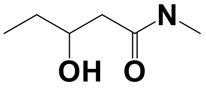 | 95 | - | - | - | - | - | - | - |
| D93 | $C_6H_6O$ | 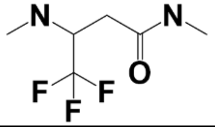 | 91 | 88 | - | - | - | - | - | - |

**D.** SAR studies exploring one side of the N'-(4-cyanophenyl)-N-aryl Malonamide

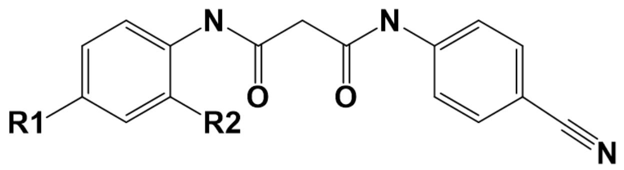


| Compound | | | % Production | | | | | | | |
|---|---|---|---|---|---|---|---|---|---|---|
| **Name** | **R₁** | **R₂** | **Pyocyanin** | ***pqsA*-GFP** | **HHQ** | **PQS** | **HQNO** | **2-AA** | **DHQ** | **AA** |
| D41 | Cl | | **6** | **8** | **2** | **9** | **15** | **6** | **11** | **739** |
| D42 | Br | | **6** | **8** | **2** | **8** | **14** | **6** | **11** | **701** |
| D43 | CN | | **8** | **8** | **22** | **24** | **39** | **19** | **34** | **531** |
| D48 | F | Me | 60 | 44 | 96 | 90 | 94 | 77 | 91 | 59 |
| D49 | COMe | | 66 | 28 | 120 | 119 | 101 | 90 | 106 | 131 |
| D50 | $COCF_3$ | | 84 | 102 | 107 | 104 | 103 | 140 | 89 | 176 |
| D51 | $NO_2$ | | 3 | 10 | 3 | 6 | 18 | 7 | 12 | 263 |
| D52 | $NH_2$ | | 87 | 90 | 92 | 86 | 92 | 101 | 103 | 115 |
| D53 | NHCOMe | | 103 | 85 | 75 | 88 | 85 | 93 | 98 | 56 |
| D54 | $NHCOCF_3$ | | 94 | 85 | 103 | 101 | 101 | 117 | 108 | 82 |
| D55 | $NHSO_2Me$ | | 100 | 131 | 130 | 115 | 98 | 95 | 80 | 117 |
| D56 | I | | 13 | 6 | 63 | 62 | 69 | 72 | 136 | 180 |
| D69 | $CF_3$ | | **4** | **2** | **6** | **16** | **40** | **15** | **13** | **920** |
| D57 | $C_6H_5O$ | | **6** | **4** | **58** | **36** | **54** | **38** | **65** | **298** |

**E.** SAR studies retaining the 4-phenoxy phenyl motif of the NAMs and exploring the effect of the substituents on the second Aryl group

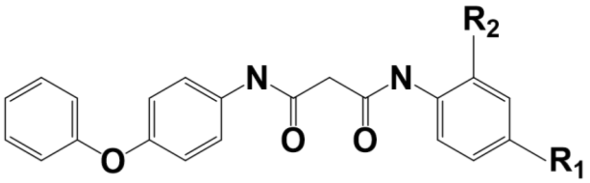


| Compound | | | % Production | | | | | | | |
|---|---|---|---|---|---|---|---|---|---|---|
| **Name** | **$R_1$** | **$R_2$** | **Pyocyanin** | ***pqsA-GFP*** | **HHQ** | **PQS** | **HQNO** | **2-AA** | **DHQ** | **AA** |
| D57 | CN | | **6** | **4** | **58** | **36** | **54** | **38** | **65** | **298** |
| D58 | NO2 | | 6 | 9 | 20 | 19 | 23 | 22 | 45 | 101 |
| D59 | I | | 72 | 81 | 99 | 93 | 97 | 92 | 93 | 131 |
| D60 | NO2 | NH2 | 6 | - | - | - | - | - | - | - |
| D61 | NO2 | OH | 6 | 7 | 1 | 6 | 8 | 12 | 8 | 112 |
| D62 | CN | NH2 | 7 | 6 | 2 | 9 | 13 | 10 | 16 | 107 |
| D63 | CN | OH | 6 | 7 | 0 | 4 | 4 | 5 | 4 | 68 |
| D64 | Br | | 48 | 61 | 88 | 100 | 88 | 78 | 85 | 50 |
| D67 | Cl | | 26 | 41 | 101 | 106 | 105 | 84 | 105 | 155 |
| D68 | F | | 33 | 52 | 121 | 137 | 121 | 84 | 109 | 127 |

**F**. Further investigation of NAMs

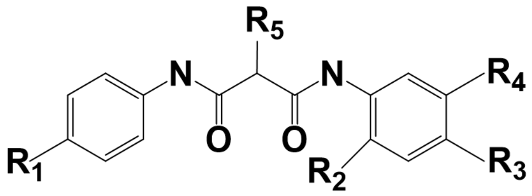

| Compound | | | | | | % Production | | | | | | | |
|---|---|---|---|---|---|---|---|---|---|---|---|---|---|
| **Name** | **R1** | **$R_2$** | **$R_3$** | **$R_4$** | **$R_5$** | **Pyocyanin** | ***pqsA*-GFP** | **HHQ** | **PQS** | **HQNO** | **2-AA** | **DHQ** | **AA** |
| D92 | $C_6H_6O$ | $NH_2$ | CN | | | 4 | 1 | - | - | - | - | - | - |
| D98 | $C_6H_6O$ | | $CF_3$ | | | 42 | 29 | - | - | - | - | - | - |
| D70 | CN | | Me | | | 50 | - | - | - | - | - | - | - |
| D71 | CN | | | Cl | | 13 | 10 | 106 | 105 | 127 | 57 | 92 | 207 |
| D72 | CN | | | F | | 30 | - | - | - | - | - | - | - |
| D75 | CN | | | CN | | 49 | - | - | - | - | - | - | - |
| D95 | CN | | CN | Cl | | **4** | **6** | **5** | **10** | **30** | **11** | **13** | **774** |
| D80 | CN | | CN | | Me | **7** | **10** | **8** | **20** | **64** | **16** | **20** | **832** |
| D88 | $C_6H_6O$ | | CN | | F | **6** | **2** | **1** | **4** | **14** | **7** | **7** | **832** |
| D96 | CN | | CN | | F | 34 | 42 | - | - | - | - | - | - |
| D100 | $C_6H_6O$ | | $CF_3$ | | F | **7** | **2** | - | - | - | - | - | - |
| D94 | CN | | CN | | Me, F | 52 | 85 | - | - | - | - | - | - |
| D97 | CN | | CN | | F, F | 94 | 95 | - | - | - | - | - | - |

**G.** NAMs analogs bearing Pyridine motif replacing one or two Aryl groups

| **Compound** | | **% Production** | | | | | | | |
|---|---|---|---|---|---|---|---|---|---|
| **Name** | | **Pyocyanin** | ***pqsA*-GFP** | **HHQ** | **PQS** | **HQNO** | **2-AA** | **DHQ** | **AA** |
| D77 | 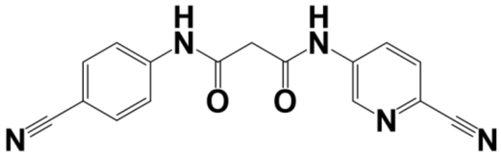 | **7** | **1** | **10** | **24** | **64** | **19** | **20** | **777** |
| D78 | 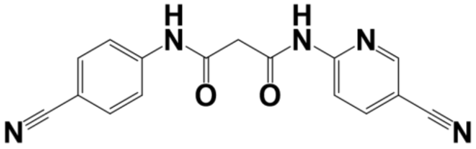 | 14 | - | - | - | - | - | - | - |
| D91 | 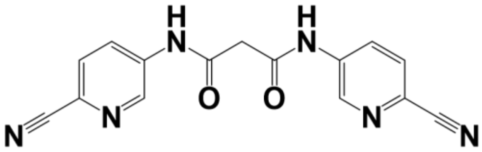 | 43 | 39 | - | - | - | - | - | - |

**Supplementary Table 2. Solubility assessment of NAMs in PBS.** Solubility of the 10mM compounds dissolved in DMSO was tested in the isotonic phosphate buffer (iPBS) at pH 7.4 using High-Performance Liquid Chromatography (HPLC). The solubility of each compound is expressed by the ratio of compound amount in the sample test solution to the amount of compound in the standard solution.

| Compound | Solubility (μM) in PBS |
|---|---|
| D41 | 13 |
| D42 | 15 |
| D43 | 11 |
| D57 | 7 |
| D69 | 21 |
| D77 | 39 |
| D80 | 29 |
| **D88** | **490** |
| D95 | 17 |
| D100 | 5 |

**Supplementary Table 3. Antibiotic-resistance profile of *P. aeruginosa* clinical isolates**

| | Antibiotic Resistance Profile | | | | | | | |
|---|---|---|---|---|---|---|---|---|
| **Strain No.** | **Amik.** | **Gent.** | **Mero.** | **Pip.** | **Tobra.** | **Cefe.** | **Aze.** | **Cip.** |
| LGR-4325 | R | R | R | R | S | R | R | R |
| LGR-4326 | R | R | R | S | S | S | S | R |
| LGR-4327 | R | R | R | R | I | R | R | R |
| LGR-4328 | S | S | S | S | S | S | S | S |
| LGR-4330 | S | S | R | S | S | S | S | R |
| LGR-4331 | S | S | S | S | S | S | S | S |
| LGR-4333 | R | R | R | R | R | R | R | R |
| LGR-4334 | R | R | S | R | R | S | S | S |
| LGR-4340 | S | S | S | S | S | S | S | S |
| LGR-4343 | R | R | R | R | R | R | S | S |
| LGR-4344 | R | R | I | R | R | R | R | S |
| LGR-4348 | S | R | R | R | R | I | S | R |
| LGR-4356 | R | R | R | R | R | R | R | R |
| LGR-4362 | R | R | R | R | R | R | R | S |
| LGR-4363 | R | R | I | R | R | R | I | S |
| LGR-4364 | R | R | R | R | R | R | I | S |
| LGR-4366 | S | R | R | R | R | R | R | R |
| LGR-4368 | S | S | S | S | S | S | S | S |

Amik. = Amikacin, Gent. = Gentamycin, Mero. = Meropenem, Pip. = Piperacin, Tobra. = Tobramycin, Cefe. = Cefepime, Aze. = Azetromycin, Cip. = Ciprofloxacin.

R = Resistant; I = intermediate; S = sensitive.

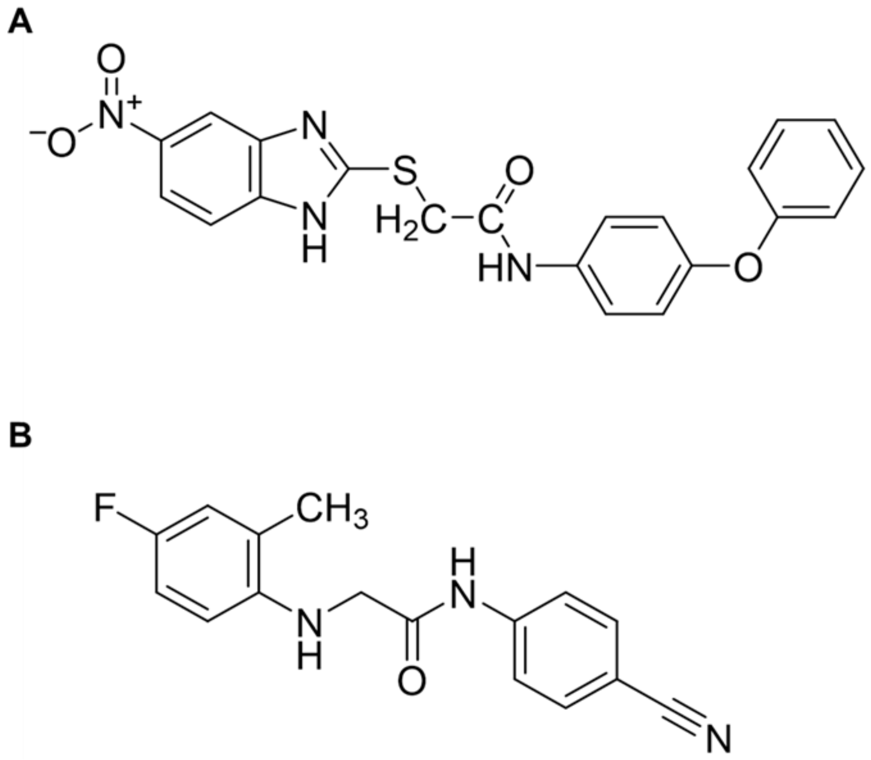


**Supplementary Figure 1. Chemical structure of (A) M64 and (B) M17**

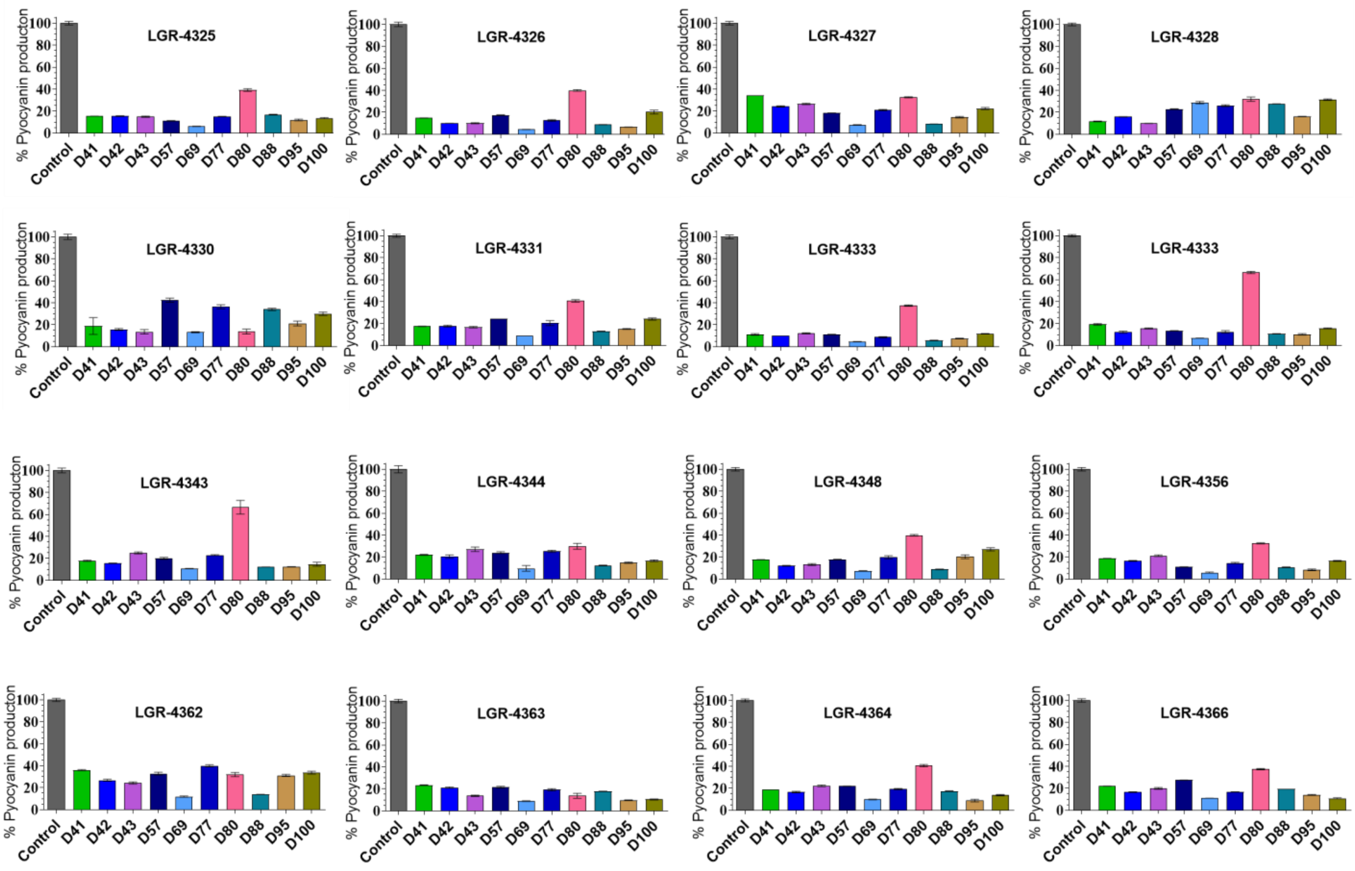


**Supplementary Figure 2. NAMs are highly efficacious against pyocyanin production in multi-drug resistant clinical strains of *P. aeruginosa*.** Pyocyanin production was measured in 16 *P. aeruginosa* clinical isolates 18hr post-growth in the presence or absence of 10μm of each NAM compound. The percent of pyocyanin production was calculated by comparing the same strain grown in the presence of the vehicle control. Data represent at least three independent replicates. The error bars denote ± SEM.

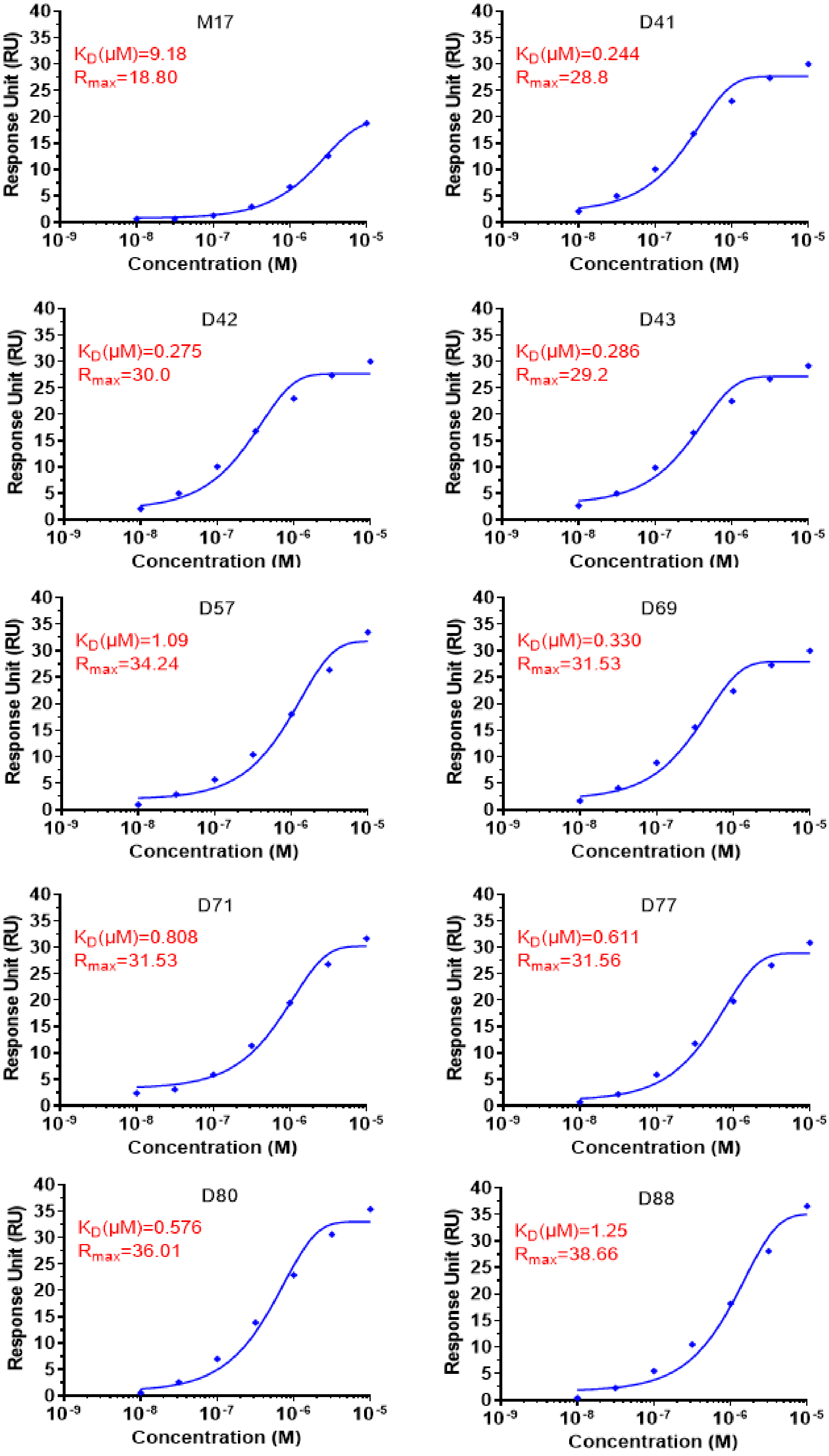


**Supplementary Figure 3. Binding kinetics profile of NAMs to the MvfR protein.** Binding of the compounds assessed using Surface Plasmon Resonance (SPR).

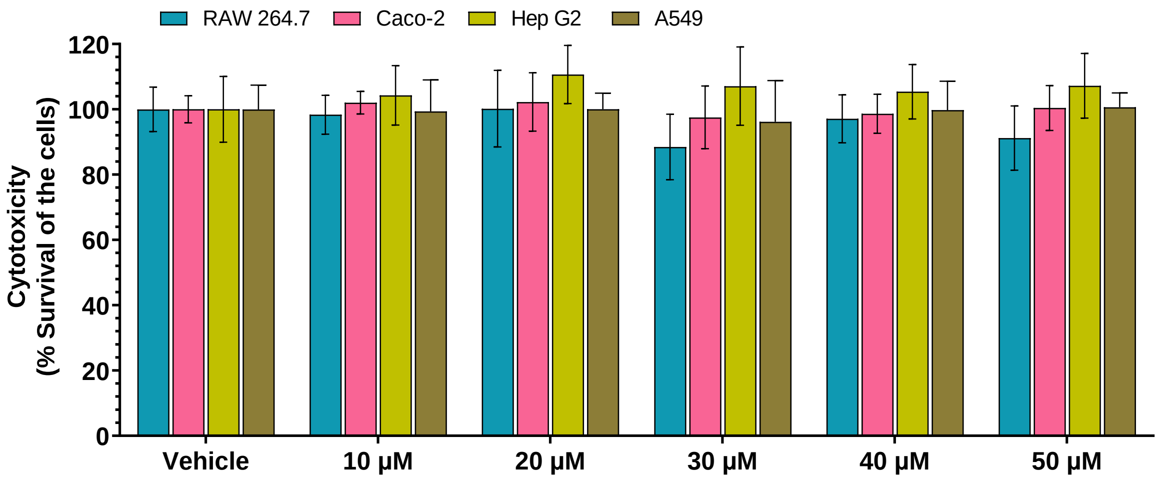


**Supplementary Figure 4. Cytotoxicity assessments of the compound D88.** Cell viability was assessed in four different cell types, RAW 264.7 (macrophage), Caco-2 (colon epithelial cells), Hep G2 (liver cells), and A549 (lung epithelial cells), the presence and absence of D88 at different compound concentrations (10, 20, 30, 40, and 50μM). None of the used compound concentrations exerted any cytotoxic effect in the tested cell lines. The cells' percent survival was calculated compared to the same cell type grown in the presence of vehicle control. Data represent at least three independent replicates. The error bars denote ± SEM.

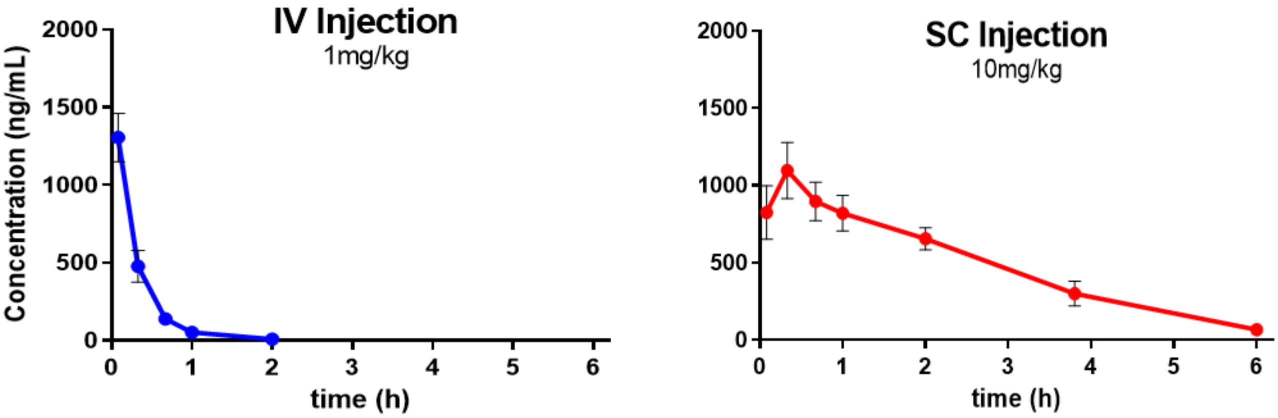


| Matrix | T ½ (h) | $T_{max}$ (h) | $T_{last}$ (h) | $C_{max}$ (ng/mL) | $C_{last}$ (ng/mL) | $AUC_{0-t}$ (h*ng/mL) |
|---|---|---|---|---|---|---|
| **IV** | 0.27 | 0.080 | 2 | 1086 | 14.2 | 494 |
| **SC** | 1.25 | 0.33 | 6 | 1096 | 66.6 | 2892 |

**Supplementary Figure 5. Pharmacokinetics of D88 *in vivo***. D88 was administered via intratracheal instillation 10.2 mg/Kg) in homogeneous suspension (0.5 % HPMC in water and assessed at different time points up to 6h.

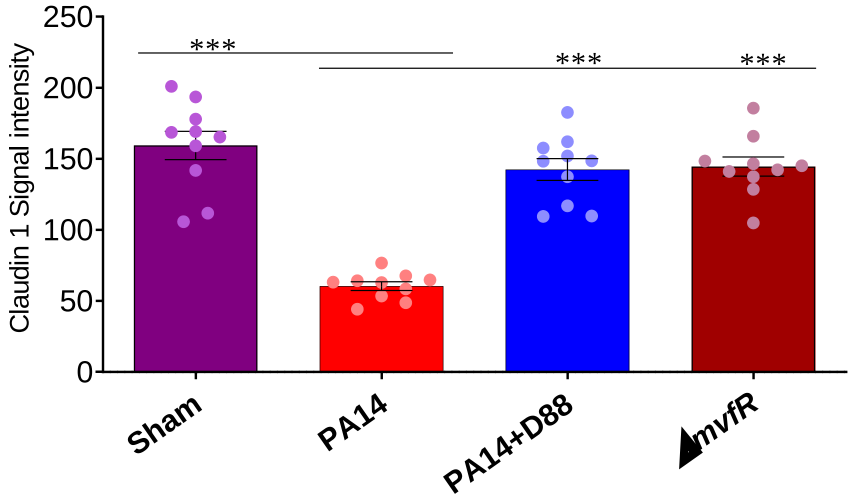


**Supplementary Figure 6. Quantification of the respective confocal images**. The signal intensity of claudin 1 in the PA14 infected + vehicle control group was compared to the groups of infected and D88-treated, *mvfR*-infected, and sham (vehicle-treated only). The error bars denote ± SEM. "***" indicate significant differences compared to the no treatment control at P < 0.001.

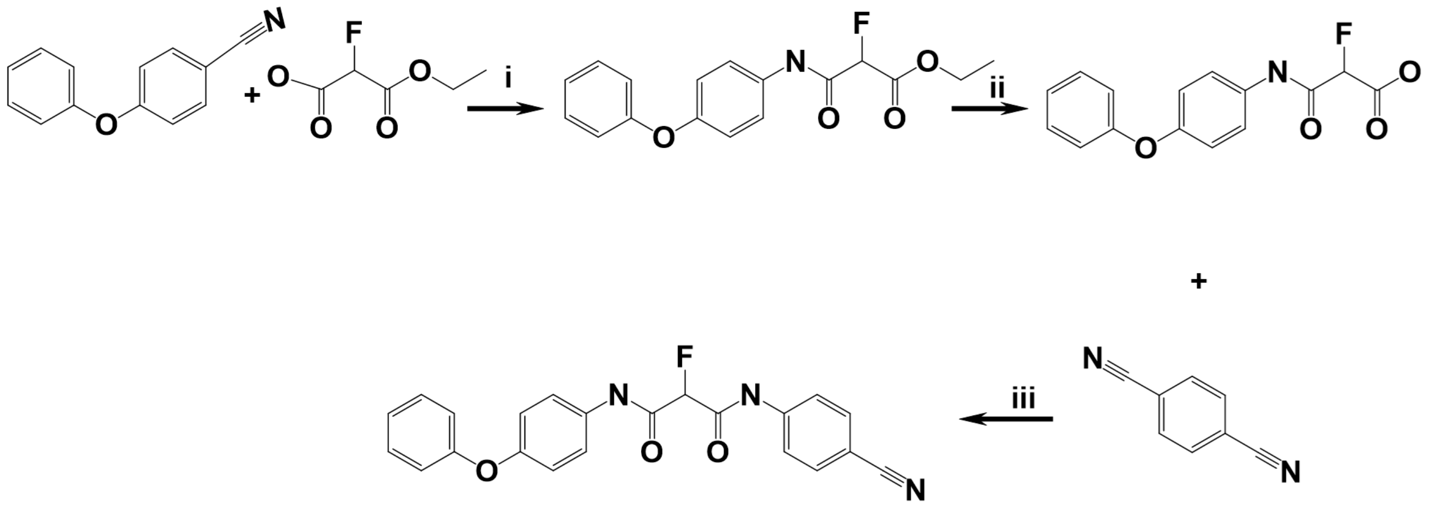


**D88**

**Supplementary Figure 7. Route of synthesis for compound D88.**
